# Tryptophan became part of the universal genetic code post-LUCA

**DOI:** 10.64898/2026.05.28.728509

**Authors:** Sawsan Wehbi, Nhan Ly-Trong, Andrew Wheeler, Bui Quang Minh, Dante Lauretta, Joanna Masel

**Author notes:** **Corresponding author:** Joanna Masel.

## Abstract

We evaluate whether tryptophan (W), widely thought to be the last of the 20 canonical amino acids added to the genetic code, was already present in the Last Universal Common Ancestor (LUCA). We reconstruct the evolutionary history of tryptophanyl-tRNA synthetase (WRS), the enzyme that attaches W to its tRNA, and the related tyrosyl-tRNA synthetase (YRS). We identify and exclude sequences derived from ancient recombination between archaeal and bacterial YRSs. Diverse rooting methods, including a novel approach exploiting time non-reversible evolution, all place the root between bacterial and archaeal YRS rather than between YRS and WRS. This supports post-LUCA WRS origination in Archaea, followed by its horizontal transfer to Bacteria. However, ancestral sequence reconstruction suggests that Archaea were depleted for W while Bacteria were not, and enzymes essential for W biosynthesis emerged in Bacteria. This suggests that W usage originated in Bacteria, with later WRS emergence in Archaea allowing the archaeal genetic code to converge with the bacterial code. The universality of the genetic code is usually attributed to common descent from LUCA, but the final step making the code universal was instead achieved by horizontal gene transfer. This gives credence to similar mechanisms for earlier steps in genetic code evolution.

## Main

The standard genetic code by which 64 nucleotide triplets are translated into 20 amino acids is claimed to be complete at the time of the Last Universal Common Ancestor (LUCA)^1,2^. The genetic code, like any complex product of evolution, must have evolved in stages, beginning with fewer amino acids and progressively adding more^3-7^. Studying these stages would be harder if they all happened before LUCA. Here we revisit whether the final amino acid, tryptophan (W)^8,9^ was already in place in LUCA. We address the incorporation of W into the genetic code by studying the origin of the catalytic domain of tryptophanyl-tRNA synthetase (WRS), which recognizes and binds to W and activates it to form Trp-AMP in preparation for its attachment to cognate tRNA. WRS evolved by duplication and divergence from its closest evolutionary relative, tyrosyl-tRNA synthetase (YRS)^10^.

Previous evolutionary analyses have not resolved whether WRS emerged before or after LUCA. The answer depends on where the root of the WRS/YRS phylogenetic tree is placed (Figure 1). Some studies supported a pre-LUCA divergence (Figure 1A)^11-14^, while others supported a post-LUCA origin (Figure 1B)^15-18^ (see Supplementary table 1). However, root support has generally been weak. Some studies assume pre-LUCA divergence as a starting constraint^19,20^.

**Figure 1.**
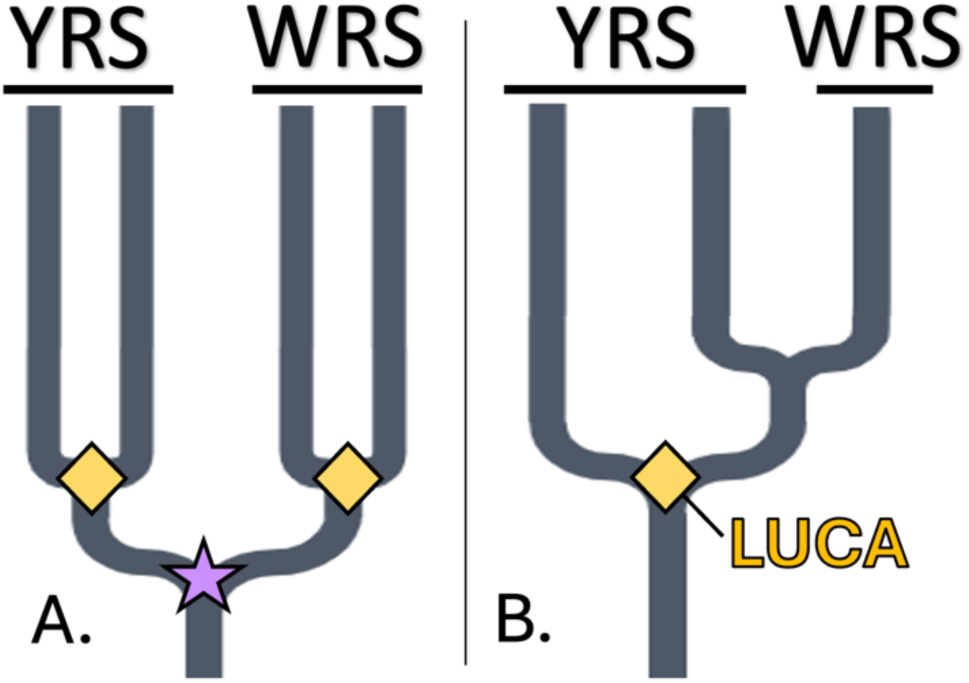
Whether WRS diverged from YRS before (left) vs. after LUCA (right) can be inferred from the root placement. LUCA (yellow rhombus) corresponds to an archaeal-bacterial split. A “pre-LUCA” duplication event is shown with a purple star.

We revisit whether LUCA had already evolved WRS. This requires new methods to overcome four main challenges. First, low sequence similarity between WRS and YRS makes ancient events hard to resolve^21^. We therefore adopt recent phylogenetic methods that exploit more slowly evolving 3D protein structures^22^. Second, Because WRS/YRS sequences are short, they provide limited information, which we supplement by reconciling^23^ our WRS/YRS gene tree with a species tree. Third, we identify and exclude an ancient recombination event between archaeal YRS and bacterial YRS, which would otherwise distort both sequence alignment and phylogenetic inference.

Finally, the extreme divergence of other synthetases from WRS/YRS makes their use as outgroups for rooting less reliable^24,25^. We therefore ensure that a variety of alternative approaches all agree on the same root. We use multiple existing methods based on branch lengths within the WRS/YRS tree. Because inaccurate models of substitution processes distort branch length and even topology^26^, we use models specific to archaea and bacteria. Our most innovative rooting approach exploits the fact that evolution in forward time is different to evolution in reverse time (e.g. methylated cytosines are subject to frequent deamination to mutate to thymine in a manner not balanced by reverse mutations). Time non-reversible substitution models exploit this asymmetry to infer a maximum likelihood root^27^.

With these improved and diverse methods, we find strong evidence that WRS emerged post-LUCA, within Archaea. In this light, we were surprised to find that ancient archaea, unlike ancient bacteria, lacked key W synthesis enzymes, and were depleted for W in ancestrally reconstructed sequences. W seems to have been first used in bacteria, then belatedly adopted by archaea through the emergence of WRS, which was later horizontally transferred to bacteria. The “universal” nature of the genetic code is not a simple consequence of common descent from LUCA, but also involved convergence of Bacteria and Archaea through horizontal gene transfer.

### Ancient YRS recombination

There are two highly diverged groups of bacterial YRS sequences^28^, which Yang et al.^29^ denote canonical and non-canonical (Supplementary Figure 1). We expected the two bacterial types to resemble each other more than either resembles archaea, and this is true for most of the protein, but in one region (Figure 2A, pink shading), non-canonical bacterial YRS sequences more closely resemble archaeal sequences. This patchwork pattern suggests an ancient recombination event, in which part of an archaeal YRS was transferred into a bacterial YRS lineage. By inspecting sequence logos (Figure 2A, top), we identified two likely recombination breakpoints that delimit this archaeal-like segment (red dashed vertical lines; Figure 2A).

**Figure 2.**
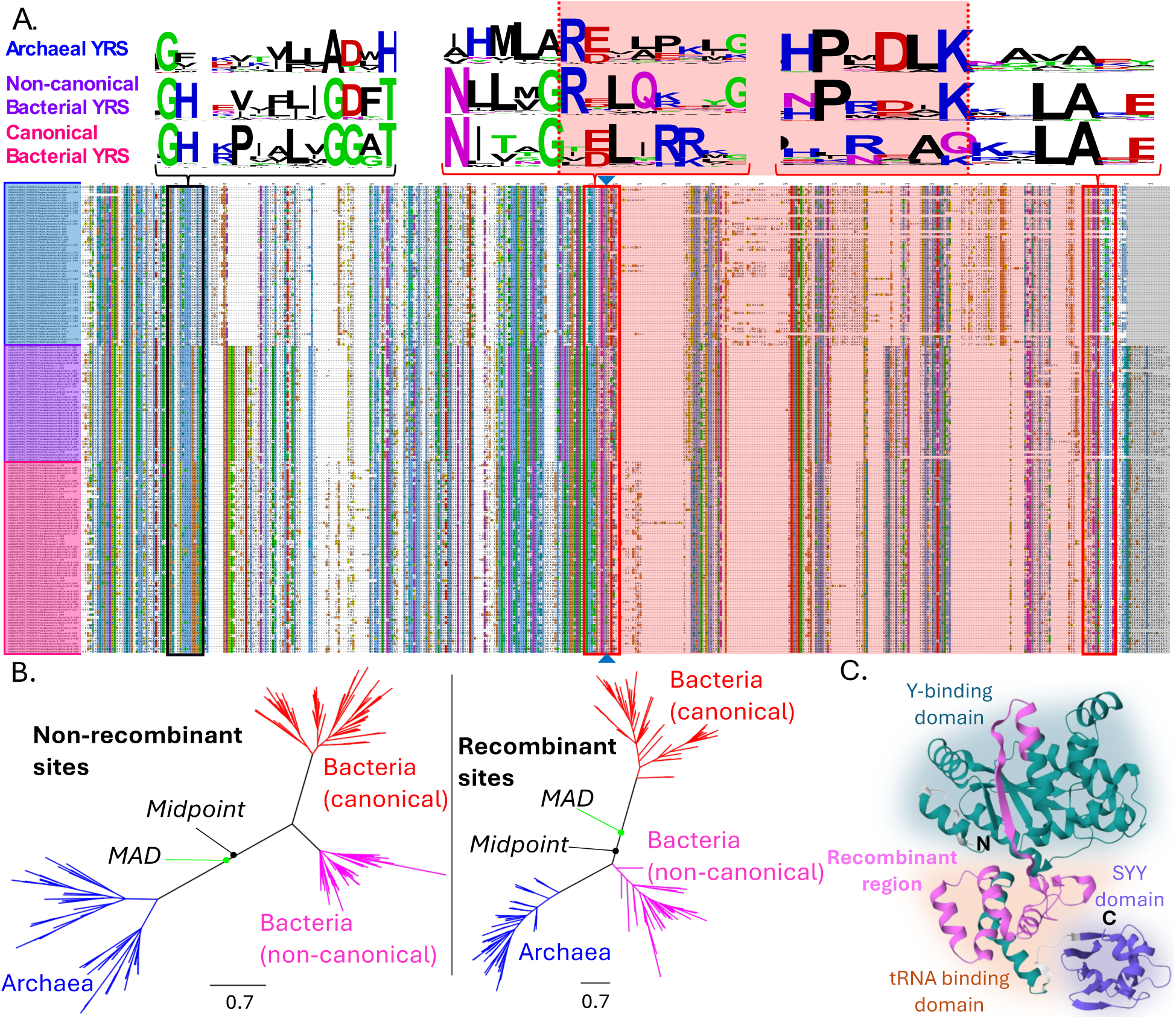
Recombination transferred part of archaeal YRS into non-canonical bacterial YRS. A) YRS core catalytic domain alignment from archaeal (indicated left as blue), bacterial non-canonical (purple) and bacterial canonical (pink) sequences. Both recombinant and non-recombinant YRSs are found within all four major bacterial supergroups (Terrabacteria, Proteobacteria, PVC and FCB) (Supplementary Figure 2). Sequence logos for the three YRS sequences types were generated using WebLogo^30^ at the three regions shown in boxes above the alignment (38-50, 209-221 and 414-425). The 448-column alignment (whose longest sequence has 319 residues) is available on sawsanwehbi/Ancient-Tryptophan-Tyrosine-usage Github repository in ‘YRS_aln.fasta’. Dashed red vertical lines indicate manually annotated recombination breakpoints. Recombinant sites are shaded in red while non-recombinant sites are left unshaded. Linker sites that contain amino acids only in bacterial YRSs have grey shading. GARD^31^ detected 3 potential upstream breakpoints with >20% support (sites 217, 295 and 296), and no downstream breakpoints. Alignment site 217 had the highest support of 38% and is indicated by triangles on the top and bottom of the alignment, 3 sites downstream of our manually annotated breakpoint. This is within the resolution of GARD, which uses alignment fragments of length 57. B) Species-tree reconciled consensus trees reconstructed for the non-recombinant (left) and recombinant YRS sites (right). Branch lengths were estimated using NQ.bac+R10 in IQ-Tree v2.1.3^32^. Outgroup rooting with six CRS sequences also places non-canonical sequences on the archaeal side of the root (Supplementary Figure 3B). C) Structure of non-canonical YRS from bacterial *Deinococcus gobiensis* (RefSeq ID WP_014686357; Uniprot ID H8GWH4). The protein sequence is 416 amino acids long and contains two annotated Pfam domains (green/pink Pfam PF00579 vs. purple S4 Pfam PF22421) and three structural modules (shown as different colored backgrounds). The white polypeptide chain is a linker region between PF00579 and the S4 domain. The recombinant sequence (pink) encompasses both the tRNA binding domain (peach background) and the Rossmann fold amino acid binding domain (green background).

Trees built separately from the two sets of sites support recombination: outside the transferred segment, non-canonical bacterial YRS groups with other bacterial YRS, whereas within the segment it groups with archaeal YRS (Figure 2B). Similarities in indel patterns between the two bacterial types within the recombinant region can be attributed to insertions in archaea; confirming this, the same sites are also absent from the cysteinyl-tRNA synthetases (CRS) outgroup (Supplementary Figure 3A). The transferred region spans parts of both the amino acid-binding and tRNA-binding portions of the enzyme and includes substantial secondary structure (alpha helices and beta sheets), indicating that the event affected functionally important parts of the protein (pink; Figure 2C).

### WRS originated in Archaea

After excluding recombinant non-canonical bacterial YRS sequences, all rooting methods placed the root between bacterial and archaeal YRS, rather than between YRS and WRS. This rooting implies that WRS did not diverge from YRS before LUCA but instead arose later within the archaeal lineage. We recovered this root both from the position of the CRS outgroup (Figure 3A), and from branch lengths on a tree topology estimated without the outgroup (Figure 3B-C). We inferred tree topologies using a partitioned model that combined information from amino acid sequence and from 3Di-encoded protein structure (see Methods), and then estimated branch lengths from one information source at a time. We used branch lengths to infer both a midpoint root and a minimal ancestral deviation (MAD) root, for both structure-based and sequence-based distances. Because the evolution of the new functionality of WRS is expected to lengthen the branch leading from YRS to WRS, the placement of midpoint and MAD roots within a long bacterial YRS branch is conservative with respect to inferring a post-LUCA origin of WRS. Support from midpoint and MAD rooting was even stronger when using the archaeal-trained amino acid substitution model NQ.arch (Supplementary Figure 4) than with the bacterial-trained model NQ.bac (Figure 3C).

**Figure 3.**
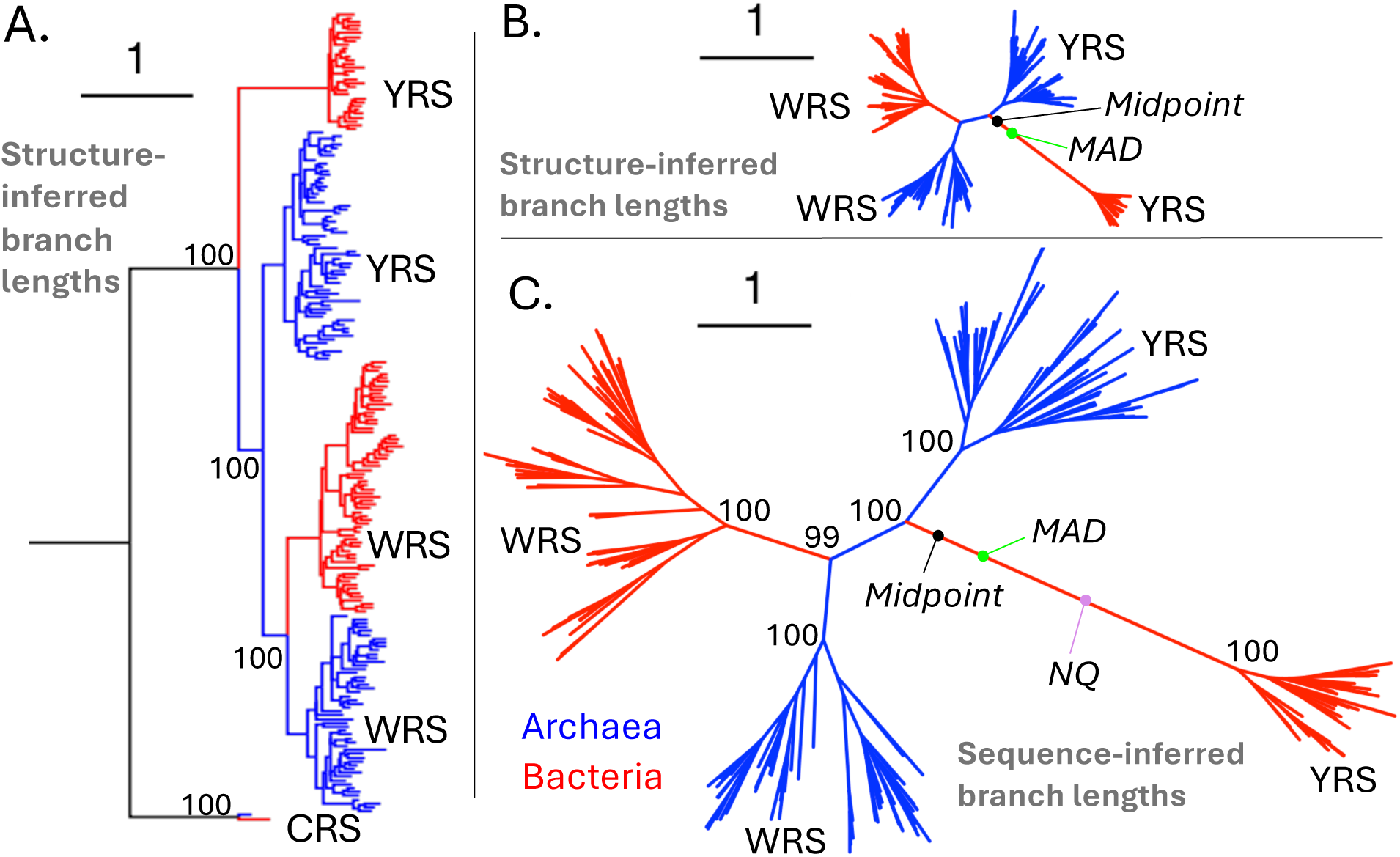
Resolved root of PF00579 domain tree reveals that WRS emerged post-LUCA from archaeal YRS. See Methods for construction of the two topologies (with and without two CRS sequences as an outgroup) from concatenation of amino acid and 3Di partitions, and reconciliation with species tree. As a visual choice in A), we placed the root halfway along the branch separating CRS’s archaeal-bacterial split from YRS. Branch lengths were inferred from A-B) Q.3Di.AF+R5, or C) NQ.bac+R10, with the time non-reversible nature of the latter producing the NQ root. The midpoint and minimal ancestral deviation (MAD) roots were estimated using midpoint() and mad() functions in the ‘phangorn’^34^ and ‘mad’^35^ R packages, respectively. Node labels in A) and C) denote bootstrap support at pertinent nodes.

Including the recombinant bacterial YRS sequences reduced the confidence of the midpoint root, placing it 0.15 amino acid substitutions (0.08% of tree length) away from the base of the bacterial YRS stem branch (Supplementary Figure 1), compared to 0.33 amino acid substitutions in Figure 3C (0.3% of tree length).

The “NQ” root, which exploit the fact that sequence divergence in forward vs. backward time have different likelihoods^27^, is even further from the base of the bacterial stem branch (Figure 3C and Supplementary Figure 4). It has 100% rootstrap support (a resampling measure analogous to bootstrap support^33^) under both the bacterial-trained and archaeal-trained time non-reversible models.

The agreement across all rooting methods supports a scenario in which YRS duplicated within the stem archaeal lineage, and then a descendant evolved into the first WRS. In this scenario, the WRS gene was later horizontally transferred into the bacterial stem lineage, prior to the divergence of bacteria into their contemporary supergroups (Figure 4A).

**Figure 4.**
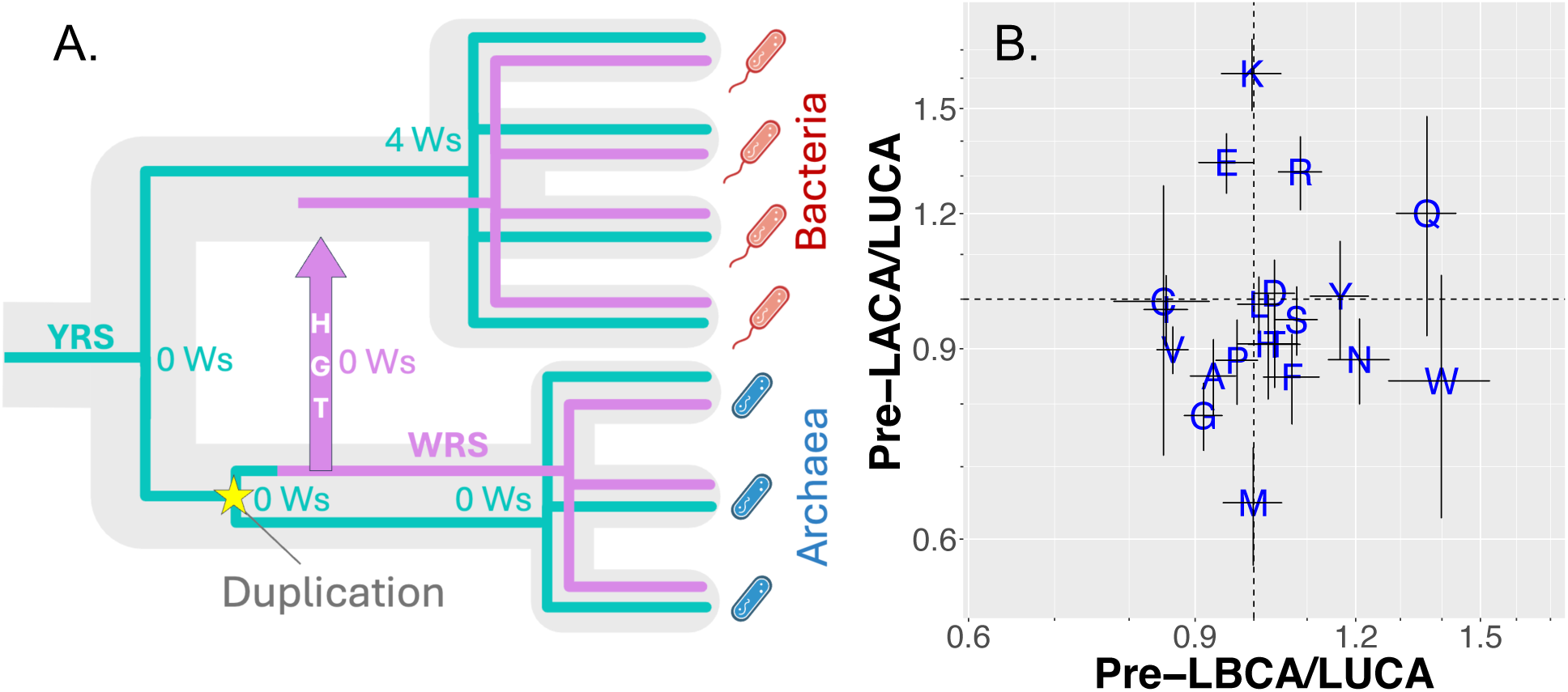
Ancient bacteria had more W sites than ancient archaea. A) The WRS (purple) and YRS (teal) catalytic domain tree, nested within the taxonomic tree. YRS duplicated in archaea (yellow star), one copy differentiated into WRS, and WRS was then horizontally transferred into bacteria. YRS contained four Ws at the last bacterial common ancestor (LBCA), but other key ancestral nodes have no high-confidence W sites (inferred with NQ.bac and shown as numbers at nodes). LBCA is shown as older than LACA^36^. B) Ancestrally reconstructed amino acid frequencies with NQ.bac+R10, in 31 pre-LACA and 251 pre-LBCA clans relative to frequencies in 309 LUCA clans (see Methods). Using NQ.arch yields similar results (Supplementary figure 6). Archaeal enrichment for positively charged arginine (R) and lysine (K) might be explained by the fact that 58% of pre-LACA vs. 11% of pre-LBCA clans are associated with negatively charged RNA/DNA (see Methods; p = 10^-6^, chi-squared test). Error bars indicate 66% confidence intervals (+/- 1 standard error) calculated on a linear scale but displayed on a back-transformed logscale, shown with tick marks at 0.1 intervals.

### Bacteria translated W before Archaea

To address whether W was present in either archaeal or bacterial proteins before the emergence of archaeal WRS, we analyzed ancestral amino acid frequencies both within the YRS/WRS family itself, and across a broader set of ancient but post-LUCA protein sequences. For the latter, we identified “pre-LACA” sequences that emerged and had already diversified within archaea prior to the last archaeal common ancestor, and “pre-LBCA” sequences that had emerged and diversified in bacteria prior to the last bacterial common ancestor (see Methods).

If archaeal WRS was the first mechanism by which W entered proteins, then ancestrally reconstructed (see Methods) pre-LACA proteins should have more W than pre-LBCA proteins do. Instead, pre-LBCA sequences have a higher ancestrally reconstructed W frequency (0.014 ± 0.0008) than pre-LACA sequences (0.008 ± 0.002), which have slightly less W even than LUCA sequences (0.010 ± 0.0006) (Figure 4B). Among all amino acids, W shows one of the strongest bacterial–archaeal discordances (Figure 4B). The four seemingly oldest archaeal sequences (see Methods) have an even lower reconstructed W frequency (0.001 ± 0.0007) across 492 ancestral sites. Newer archaeal proteins, emerging long after WRS, were also born with less W than newer bacterial proteins (Supplementary figure 5A-B). Note that the magnitude of amino acid frequency deviations from current usage will tend to be underestimated, because ancestral sequence reconstruction uses a model whose equilibrium corresponds to contemporary frequencies of the 20 amino acids (Wong et al., unpublished results).

We next assessed whether data were consistent with a pre-LACA W usage as low as zero. Around 35% of pre-LACA W comes from sites reconstructed with <70% confidence, compared to only 13% for pre-LBCA W (Supplementary figure 7A-B). This asymmetry is not seen for other amino acids such as Y (Supplementary figure 7C-D). Even high-confidence sites could be the product of convergent evolution toward a functionally advantageous W once W became available. We therefore focused on “ultra-conserved” sites at which an amino acid was conserved in 100% of sequences in our alignment (Supplementary figure 7E, top values). LBCA and pre-LBCA have 35 sites ultra-conserved for W versus 824 for other amino acids, but LACA and pre-LACA have only 1 for W versus 229 for other amino acids (p = 0.003, Fisher’s exact test). LACA’s only ultra-conserved W site is in a protein domain with unknown function (PF18898), the inference of whose history might be complicated by its presence in many viruses. These results are thus compatible with, without being proof of, the complete absence of W from early archaea, and by extension from LUCA. There are no ultra-conserved W sites in either LUCA or pre-LUCA, even though 97 sites are ultra-conserved for some other amino acid (Supplementary figure 7E).

Specifically in the YRS/WRS family, we found no strongly supported W sites in its most recent common ancestor (MRCA), consistent with Fournier and Alm^19^. Nor did we find confidently reconstructed W sites in the MRCA of archaeal YRS. In contrast, the MRCA of bacterial YRS contains four W sites reconstructed with >99% probability (Supplementary table 1; Figure 4A). At the root, while W is the single most likely residue at all of these four sites (plus one additional site when using NQ.arch), its confidence is <50% (Supplementary table 2). Ample statistical power is demonstrated by reconstructing Y with >70% confidence at deep nodes: three sites at the root, six in the YRS/WRS MRCA, and nine and twelve in the bacterial and archaeal YRS MRCAs, respectively.

### Bacteria were the first to synthesize W

To ask when the modern pathway for tryptophan biosynthesis became complete, we examined the evolutionary ages of the protein activities required for the pathway shown in Figure 5. We identify these by means of the Pfam IDs^37^ associated with that portion (domain) of the protein; different organisms have different architectures by which Pfams are mixed and matched into complete proteins. Two essential domains, PF00218 and PF00425, appear only in the bacterial lineage, implying that the full pathway was completed in bacteria rather than in LUCA.

**Figure 5.**
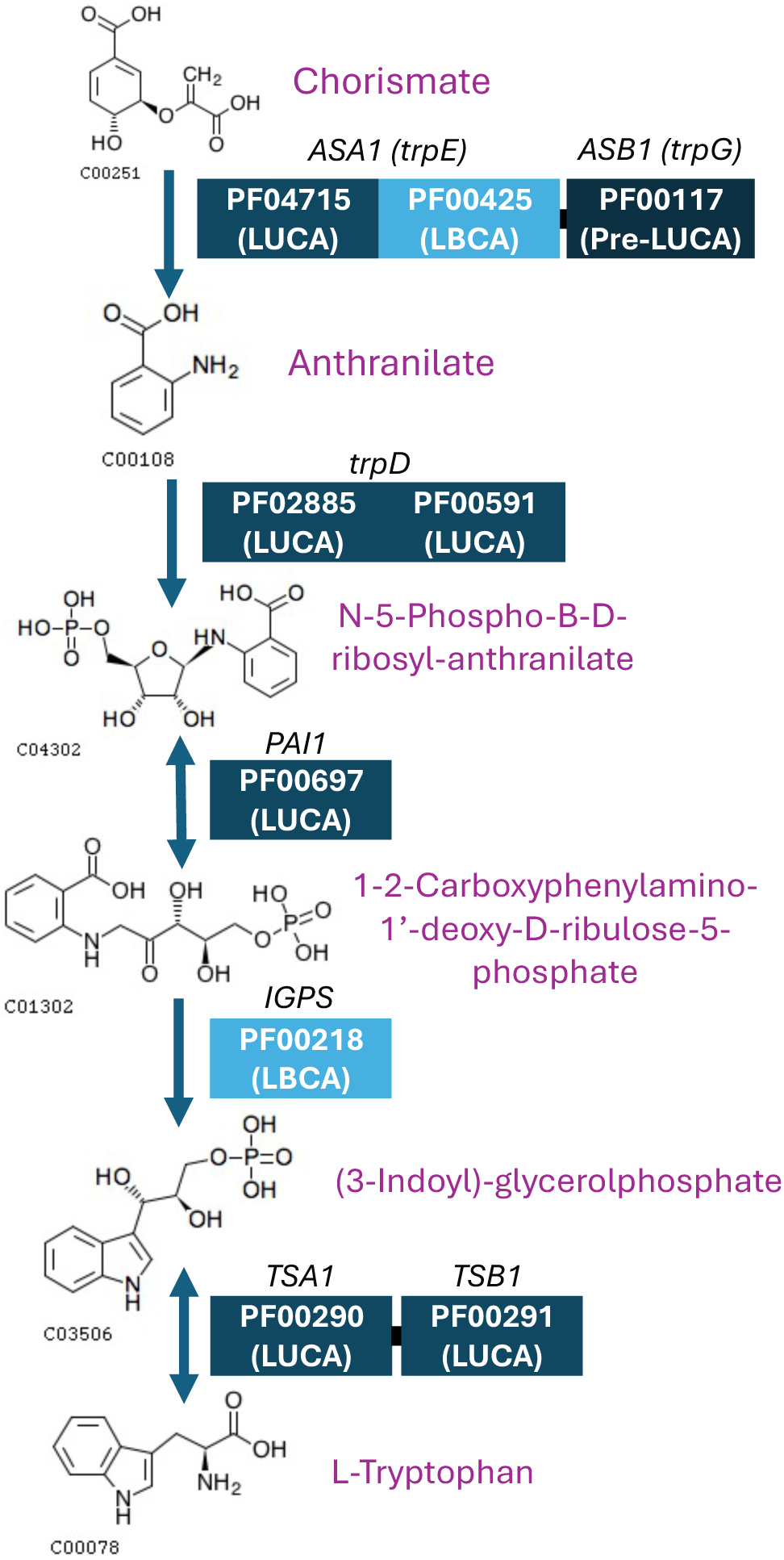
Tryptophan synthesis pathway from Radwanski and Last^43^. Presentation is adapted from KEGG tryptophan biosynthesis pathway (M00023)^42^. Compound names are shown in purple next to their respective structures. Boxes represent Pfam domains of enzymes, with the Pfam IDs and their classified ages from Wehbi et al.^8^ in white. Gene names are shown in black, above the boxes. Thick black lines connect two separate genes (ASA1 and ASB1) that form the anthranilate synthetase complex, and connect the tryptophan synthetase alpha and beta chains (TSA1 and TSB1). We show one possible arrangement of domains into genes; some domains are sometimes found fused with other components in the pathway, giving rise to bifunctional proteins. For example, a tryptophan biosynthesis protein now known as TrpCF contains both PF00697 and PF00218^38^.

PF00218 is an Indole-3-glycerol phosphate synthase (IGPS) domain that catalyzes a key ring-closing step in W biosynthesis^38^. Consistent with our results, previous studies did not infer the corresponding gene IGPS (K01609) to be part of LUCA’s proteome^39,40^.

PF00425 is the part of the anthranilate synthase that binds chorismate, the starting molecule for this step in W biosynthesis^41^. One recent study inferred that the corresponding gene trpE (K01657) was 50% likely present in LUCA^40^. However, that inference may reflect the presence of its other domain (PF04715), which traces back to LUCA^8^, while the chorismate-binding domain itself, PF00425, does not. Because anthranilate cannot be synthesized from chorismate without this binding activity, our results suggest that this step of the modern W biosynthesis pathway was not yet complete in LUCA.

Archaea now widely engage in tryptophan biosynthesis, following horizontal gene transfer of the pathway from Bacteria. In the KEGG database^42^, 76% of 473 sampled archaeal genomes contain IGPS (K01609), 79% contain trpE (K01657), and 70% contain the complete modern W biosynthesis pathway (M00023). This complete pathway is distributed across all four major archaeal supergroups.

## Discussion

A broad variety of tree rooting methods all support a post-LUCA, archaeal origin of WRS, after implementing recent innovations in structural phylogenetics, time non-reversible models, and species tree reconciliation. The evidence becomes stronger when an extraordinarily ancient recombination event between bacteria and archaea is accounted for. Despite the archaeal origin of WRS, ancient bacteria used more W than ancient archaea, and indeed ancient archaea might not have used W at all. This is supported by sequence reconstruction both of WRS/YRS ancestors, and more broadly of other ancient protein domains. It is also supported by the fact that the W biosynthetic pathway was completed in Bacteria, and was absent from LUCA.

We are not aware of any previously reported recombination events this ancient, predating the last common ancestors of both Archaea and Bacteria. Such an event is nonetheless plausible because horizontal transfer was common for ancient aaRSs^28,44,45^, as well as more recently for YRS in particular^46^, creating opportunities for recombination. A more recent recombination between two different supergroups of archaea also involved YRS; Crenarchaeota donated a segment to YRS in a subset of Halobacteriales^19^. Distantly related forms of other aminoacyl-tRNA synthetases have also recombined. A single archaeal species, *Natrialba magadii,* acquired a bacterial N-terminal LeuRS domain, in combination with an otherwise archaeal LeuRS^47^. A recombinant proteobacterial GlxRS has unknown origins for both anticodon- and amino acid- binding portions^48^. ThrRS from two crenarchaeotal genera (*Sulfolobus* and *Aeropyrum*) contain a stretch of 150 residues from an unknown donor^28^. Fournier et al.^19^ suggested that successful recombination is constrained by the need to preserve coevolution between the enzyme and its cognate tRNA. The recombination we found apparently overcame this constraint, with an archaeal anticodon-binding domain, including the archaeal version of the KMSKS motif that is crucial for tRNA binding, being successfully transferred into a bacterial lineage^49^. Another exception is a WRS sequence in the bacterium *D. radiodurans* (Phylum Deinococcota) whose C-terminal resembles that from the bacterial Phylum Aquificota^28^.

A plausible timeline for the incorporation of W into the translation machinery would be that 1) LUCA had only 19 amino acids, then 2) stem bacteria biosynthesized W and translated W via an unknown mechanism, then 3) the complete W biosynthesis pathway was horizontally transferred to stem archaea, which rapidly evolved WRS or acquired it from a now-extinct “ghost lineage”^50^, followed by 4) transfer, with displacement, of WRS from archaea to bacteria. We note that horizontal transfer of archaeal synthetases to Bacteria is more common than the reverse^28^. Stem bacteria might have acquired W from their environment, before they evolved the ability to synthesize it. Indeed, alkaline hydrothermal vents abiotically produce W^51^, consistent with LBCA being a hyperthermophile^52^.

If one looked only at the WRS tree, one would incorrectly presume that its root represented LUCA’s speciation into archaea and bacteria rather than horizontal gene transfer. This raises the question of how many other apparent LUCA nodes are products of unidentified horizontal gene transfer. An extreme version of this view is that all inferred LUCA nodes represent horizontal transfer rather than vertical speciation events.

Vetsigian et al.^2^ proposed that multiple forms of early life converged on a universal genetic code, driven by network effects^53^. Prior to this, life tolerated enormous ambiguity in order to permit any horizontal gene transfer. Convergence on the universal genetic code enabled both more effective horizontal gene transfer of innovations, and more refined adaptation under an unambiguous code. We note that a burst of transfer between archaeal and bacterial cells, as defined by their distinct membrane chemistries, could have given rise to the various LUCA gene tree nodes. This view helps explain why archaeal versus bacterial membranes have lipids of opposite chirality and different chemistry^54^.

Our findings align with a modified version of the Vetsigian et al. hypothesis^2^, where code convergence was not yet complete at the time of LUCA. Specifically, bacteria already had W, while archaea did not. The last stage of convergence took surprisingly long, such that we were able to root the WRS/YRS tree. Paradoxically, the incomplete nature of the genetic code at the time of LUCA is what might have preserved the evidence that now allows us to reconstruct how convergence on W usage occurred.

Bacteria and archaea might have converged in other ways too. For example, the extreme depletion of ancient but not modern archaea for methionine (Figures 4B and Supplementary Figure 5) suggests that ancient archaea might have also lacked its derivative/precursor, S-adenosyl-methionine (SAM), and that SAM-dependent metabolism might therefore be derived from the bacterial pre-LUCA ancestor. LUCA and pre-LUCA enrichment for methionine might indicate a high rate of transfer of methionine-rich proteins from ancient bacteria to ancient archaea.

While the “universal” genetic code is not in fact used by all life, even its variants are generally believed to stem from common descent with modification from LUCA^55^. In contrast, under the hypothesis of Vetsigian et al.^2^, the genetic code of one lineage might have adopted an amino acid before another lineage did. Here we found evidence that this code convergence hypothesis is correct in the case of W. While we cannot say for certain how the other 19 standard amino acids came to be shared and encoded by the same codons, our results make a broader version of the code convergence hypothesis more plausible. Under this view, the universal genetic code might have arisen through the integration of innovations developed independently by different lineages, some of them perhaps now extinct, rather than arising by a series of temporally constrained steps within a single vertically inherited lineage.

## Methods

### Archaeal amino acid substitution model

We used a previously reported bacteria-specific amino acid substitution model NQ.bac^56^, and produced a new NQ.arch model for archaea. We downloaded alignments of archaeal protein sequences from the HAMAP (High-quality Automated and Manual Annotation of Proteins) database^57^ in November 2024. We took a total of 688 alignments from the Archaea-only, Archaea+Bacteria and Archaea+Bacteria+Eukaryota datasets. We kept only the archaeal sequences from the two latter datasets. We re-aligned with Muscle5^58^. The alignments had on average 171 archaeal species (with exactly one sequence per species), and 822 sites representing the alignment of 299 amino acid residues. We estimated a time non-reversible model from the alignments using the nQMaker functionality of IQ-Tree v2.1.3^27^ with the option --model-joint NONREV+F0^59^. The archaeal-trained time non-reversible model (NQ.arch) and the archaeal alignments it was trained on can be found on the sawsanwehbi/Ancient-Tryptophan-Tyrosine-usage GitHub repository.

### WRS/YRS amino acid PF00579 sequences

YRS and WRS share both an amino acid binding and a tRNA binding domain, which while structurally modular, are annotated as a single Pfam domain (PF00579). We began with the 908 sequences of the WRS/YRS core catalytic domain (PF00579) from Wehbi et al.^8^. We exclude the additional S4 C-terminal domain (either PF01479 or PF22421), which mediates RNA binding in other proteins^60^. The S4 domain is found in most bacterial YRS (Supplementary Figure 8; no strong phylogenetic pattern) and would inflate the distance of bacterial YRSs from other PF00579 genes.

We downsampled from 908 sequences to 356 while preserving phylogenetic diversity. We began with one randomly sampled sequence out of the 908 and sequentially added sequences that added diversity. We calculated tip-to-tip distances from branch lengths inferred by Wehbi et al.^8^ with NQ.Pfam^27^ for their species-tree reconciled PF00579 tree. In each step, we considered 10 candidate sequences, and chose the one whose minimum distance to any of the existing set was largest. We repeated the process until we selected 400 sequences. We then removed all 43 sequences from the CPR bacterial supergroup due to their ambiguous evolutionary history^61,62^. We also removed a truncated archaeal sequence belonging to an Euryarchaeota genome ‘G000403645’, bringing the total to 356 PF00579 sequences: 122 for bacterial WRS, 56 for archaeal WRS, 117 for bacterial YRS, and 61 for archaeal YRS.

### Recombination identification

We first aligned all 117 bacterial and 61 archaeal PF00579 YRS sequences using Muscle5^58^. We distinguish 44 “non-canonical” bacterial YRS sequences that are deeply diverged from 73 “canonical” bacterial YRS sequences, where the longer branch leads to the canonical group (Supplementary Figure 1). We confirm that non-canonical YRS have a lysine residue at the tyrosine binding site (alignment site 9 in ‘YRS_aln.fasta’) instead of the Y residue found in all the canonical and archaeal YRSs^12^.

On the basis of visual inspection of position weight matrices, we divided the YRS alignment into two sets of columns: the putative recombinant sites (214-419), and the nonrecombinant sites (1-213 and 420-430). We also ran Genetic Algorithm for Recombination Detection (GARD) to confirm the presence of recombination and the approximate location of the breakpoint^31^. Due to the computational limitation of the GARD webserver, we took only 30 YRS aligned sequences by randomly sampling ten from each of the bacterial groups and archaea. We truncated the alignment at site 430, excluding flanking C-terminal sites unique to bacterial YRS. We set the site rate heterogeneity to general discrete with two rate categories.

To ensure that the partial horizontal gene transfer in the non-canonical YRS was not acquired from a non PF00579 sequence, we used the HMMER webserver^63^ to build a hidden Markov model (HMM) from the alignment of the recombinant non-canonical bacterial YRS sites. We then used the recombinant YRS HMM to search the reference proteomes sequence database (2025_01) from UniProt^64^. The top hits corresponded to YRS followed by WRS sequences, confirming that the recombinant region arose from YRS.

We then used Muscle5^58^ to align the recombinant portion of YRS sequences with homologous portions from six outgroup cysteinyl-tRNA synthetase (CRS): three archaeal (one from each Euryarchaeota, TACK and Asgard), and three bacterial (one from each Proteobacteria, Terrabacteria and PVC). We chose CRS as an outgroup since we are not aware of reported instances of recombination, unlike GluRS^48^ and LeuRS^47^. CRS aligned satisfactorily.

We used ModelFinder in IQ-Tree^65^ to select the R10 rate heterogeneity model as the best fit for both recombinant (both with and without CRS) and non-recombinant YRS alignment subsets (without CRS), as manually annotated. For each subset, we generated two tree ensembles, using the Bacteria-trained time non-reversible model (NQ.bac) +R10^56^, and using NQ.arch +R10, from the ultrafast bootstrap option -B 1000^66^. We used each of these as a prior distribution in AleRax^67^, to reconcile with the Web of Life (WoL) prokaryotic species tree^68^, using the UndatedDTL reconciliation model and a posterior gene tree sample set of 500. We merged the reconciled NQ.bac and NQ.arch posterior tree distributions, then unrooted the resulting ensemble of 1000 reconciled trees using the unroot() function in the ‘ape’ R package^69^. We inferred a consensus tree across the merged posterior set, separately for the recombinant and non-recombinant subsets, using the -con option with 100 burn-in and 0.5 minimum threshold in IQ-Tree v2.1.3^32^.

### Amino acid and 3Di partitioned alignment

After removing the 44 recombinant non-canonical bacterial YRS sequences, we mapped the remaining 312 PF00579 sequences to their Uniprot IDs, 209 of which had structures available on AlphafoldDB^70^. We downloaded the 209 AlphaFold structures as PDB files. We converted them into amino acid residues and corresponding 3Di characters using the Foldseek webserver^71^.

Re-alignment after removing recombinant sequences is crucial. When recombinant sequences are present, a single guide tree is imposed across all sites during multiple sequence alignment, even though different regions follow distinct evolutionary histories. As a result, the alignment process attempts to reconcile conflicting phylogenetic signals, particularly within recombinant segments. Thus, even if recombinant sequences are removed after alignment, the alignment itself may already reflect these inconsistencies.

We first constructed an alignment with FoldMason^72^ in order to use information from 3Di-encoded structure as well as sequence, but it had a misaligned ‘KMSKS’ motif that was clearly incorrect in the sequence alignment. We therefore aligned the 209 amino acid YRS and WRS sequences with Muscle5^58^, and imposed this Muscle5 alignment onto our 3Di sequences.

Alignments often have regions unique to only one group in the dataset, especially in the flanking regions. Likelihood calculations treat the corresponding gaps as an equilibrium amino acid frequency distribution, which can create artifacts in downstream trees. We therefore extracted just the shared PF00579 domain from the full-length alignments obtained above, using its coordinates from InterPro^73^ within a WRS protein sequence (I3IM11) (positions 4 to 287 in the 333 amino acid long protein). Linker sequences from PF00579 to the S4 domain can be seen as a C-terminal flanking region in YRS in Figure 2A, where the S4 C-terminal domain was excluded but the linker remained. This region was trimmed on the basis of absence from I3IM11 WRS.

Finally, we concatenated the amino acid and 3Di alignments, treating them as two partitions. The trimmed and partitioned AA+3Di alignment is available in ‘TruncPF00579_AA3Di_Musclealn.fasta’ on GitHub.

We included two full CRS proteins, one from bacteria (Q06752) and one from archaea (Q8U227), as an outgroup. We downloaded their AlphaFold structures as PDB files and converted them into amino acid and 3Di characters in Foldseek^71^. We re-aligned the two full-length CRS amino acid sequences with the 209 WRS and non-recombinant YRS, using Muscle5 ^58^, imposing this alignment also onto our 3Di sequences.

For both the amino acid and 3Di alignments, we again used the coordinates of PF00579 in the I3IM11 WRS to extract PF00579 together with its homologous region in CRS. We trimmed 101 and 109 amino acid residues from bacterial and archaeal CRS, respectively, because they had no homologous residues in WRS or YRS. We found only two alignment sites not present in CRS, which we did not trim. Including the CRS outgroup to the alignment introduced a few more gaps, lengthening the alignment from 741 without CRS to 755 columns. The trimmed and partitioned AA+3Di alignment with the CRS outgroup is available in ‘Trimmed_TruncPF00579_AA3Di_MusclealnCysRS.fasta’ on GitHub.

### Structure-informed phylogenetic trees

We inferred 1000 ultrafast bootstrap trees with -B 1000 and -Q in IQ-Tree v2.1.3^32^ for the two AA+3Di partitioned alignments, one with and one without CRS. We assigned NQ.bac^56^ + R10 and Q.3Di.AF^74^ +R5 for each partition, respectively^75^. We also perform an alternative analysis with NQ.arch + R10, instead of NQ.bac + R10, for the amino acid partition, in the analysis without CRS.

Partition models combining amino acids with 3Di characters can improve tree resolution for deep divergences^76^, but failed to outperform sequence-based maximum likelihood methods with less diverged phylogenomic datasets^77^. These two studies assumed an edge-proportional partition model where branch lengths for the amino acid sequences are proportional to corresponding branch lengths for the 3Di sequences (option -p in IQ-Tree). However, assuming proportional rates between partitions^78^ can produce bias and statistical inconsistency^79^. Garg and Hochberg^74^ therefore argued that AA+3Di partition models likely require edge-unlinked models^75^ that allow each partition to have its own, independent set of branch lengths. Indeed, when we tried to fit the edge-proportional model, it failed, triggering a warning that a scaling factor, describing the ratio of branch lengths across partitions, was too high.

We reconciled each ensemble of bootstrapped gene trees with the WoL prokaryotic species tree^68^ using AleRax^67^, setting the --gene-tree-rooting option to ROOTED. We then inferred a consensus tree in IQ-Tree v2.1.3^32^ using the -con option with 100 burn-in and 0.5 minimum threshold.

Accurate branch lengths are critical to root inference without an outgroup. For the consensus tree without CRS, we re-estimated branch lengths once by applying NQ.bac + R10 to the amino acid sequences, once by applying NQ.arch+ R10, and once by applying Q.3Di.AF+R5 to the 3Di sequences. Use of NQ.bac or NQ.arch enables root inference from the fact that likelihoods are not equal in forward versus reverse time^27^. We calculated associated rootstrap support values using the species-tree reconciled ensemble of trees with the --rootstrap option in IQ-Tree v2.1.3^32^, while holding the tree topology constant and omitting the 3Di partition. For the consensus tree with CRS, given the extreme sequence divergence of CRS from WRS and YRS, we re-estimated branch lengths only using the 3Di sequences with Q.3Di.AF+R5.

Midpoint and minimal ancestral deviation (MAD) roots were estimated using midpoint() and mad() in the ‘phangorn’^34^ and ‘MAD’^35^ R packages, respectively. Wade et al.^80^ found the MAD root to be superior than midpoint, based on the “root balance” of gene trees (the ratio between the numbers of species on either side of the root) being closer to that of corresponding species trees.

### Pfam classification

Wehbi et al.^8^ classified 285 and 2770 ancient but post-LUCA Pfam domains that date back to the last archaeal common ancestor (LACA), and to the last bacterial common ancestor (LBCA), respectively. From among these, we identified Pfams for which both sides of the most basal split qualify as LACA or LBCA according to the phylogenetic trees inferred in Wehbi et al.^8^. This yields 63 ‘pre-LACA’ and 575 ‘pre-LBCA’ Pfams.

Pfams represent sequences similar enough to align, and to be represented by a Hidden Markov Model; more distant homologs are represented in the Pfam database as ‘clans’ that can contain multiple Pfams. The age of a sequence is best captured by the age of its clan, not its Pfam. Wehbi et al.^8^ identified 1232 clans, i.e. sets of Pfams classified as homologous, which date back to either LACA or LBCA, but not to LUCA. We similarly reclassify a subset of these as pre-LACA and pre-LBCA clans, for use in the numerator of ancient archaeal and bacterial amino acid usage. Pre-LACA clans include either one pre-LACA Pfam or at least two LACA Pfams. Similarly, pre-LBCA clans include either one pre-LBCA Pfam or at least two LBCA Pfams.

As control sequences, some of our analyses use LUCA clans (i.e. which predate the archaea vs. bacteria split), and others use more recently emerged “modern” clans. Wehbi et al. ^8^ classified 309 LUCA clans, requiring the presence either of exactly one LUCA Pfam (with an archaeal-bacterial split at the root), or of a LACA Pfam and a LBCA Pfam within the same clan. Wehbi et al.^8^ also classified 2234 modern clans, for which all Pfams emerged in a single prokaryotic supergroup, using previously classified supergroups^61,81-83^. Modern archaeal clans include 2 Asgard, 133 Euryarchaeota, 24 TACK, and no DPANN. Modern bacterial clans include 609 Proteobacteria, 742 Terrabacteria, 39 PVC, 120 FCB, and 4 CPR. We consider only these supergroup specific clans as modern. We exclude 501 clans that originated at the common ancestor of only two of the five bacterial supergroups, previously referred to as modern ‘post-LBCA’ in Wehbi et al.^8^. We also exclude four clans classified by Wehbi et al.^8^ as modern that contained modern Pfams belonging to both bacterial and archaeal supergroups.

To score nucleic acid association of pre-LBCA and pre-LACA clans, we matched their clan IDs (or Pfam IDs if they are single-entry clans) with their associated function using the ‘Pfam_A.hmm.dat’ file from the InterPro database^73^. We scored as nucleic acid associated the clans whose functional descriptions included “RNA”, “DNA”, or “ribo”.

### Amino acid usage

We used a developmental version of IQ-Tree (v3.0.1.esr.3) to perform ancestral sequence reconstruction^84^ that incorporates indels as possible ancestral states, using a newly implemented - gap-asr option, a version of which was used in^85^. Briefly, this option first performs ancestral sequence reconstruction to estimate posterior probabilities for all 20 amino acids at all sites in the multiple sequence alignment. It then recodes the alignment as a binary, with 0 representing gaps (indels) and 1 representing amino acids. It then repeats ASR on this recoded alignment using the best-fit binary model with branch lengths re-estimated on the fixed topology inferred in the first step. We used the results of the binary model to exclude ancestral sites inferred to have an indel probability above 30%, and then for the retained sites, used the original posterior probabilities for the 20 amino acids.

We applied this procedure to pre-LACA, pre-LBCA, modern archaea, modern bacteria, and LUCA clans, using Pfam alignments and phylogenetic trees from Wehbi et al.^8^, and using either NQ.arch+R10 or NQ.bac+R10 as the substitution and rate heterogeneity models^32^. We then excluded sites where the amino acid with the highest probability has an ancestral probability <0.35 for sequences within pre-LBCA clans, and <0.4 for all other clans. We chose these slightly different exclusion criteria to ensure that the concatenated ancestral sequence length (without gap characters) of all five age groups (LUCA, pre-LACA, pre-LBCA, modern bacteria and modern archaea), following both our exclusion steps, fell by the same amount, either ∼ 14% (NQ.bac) or ∼ 15% (NQ.arch) of the concatenated median length of contemporary sequences. Our primary “amino acid usage” metrics are ratios of ancestrally reconstructed amino acid frequencies in two different age classes. Bias in which amino acids tend to be excluded by our quality filters is thus expected to have equal effects on the numerator and denominator of our usage ratios, and hence to cancel out.

At the root of each Pfam tree, we calculated the ancestral amino acid frequencies by taking the average of the amino acid probability distributions (with indels re-assigned probability of zero) across those sites that passed our filters. For each clan, we took the average across its Pfams, weighted by the number of ancestral amino acid sites retained past our filters. For each age cohort (pre-LACA, pre-LBCA, modern archaea, modern bacteria, and LUCA), we then took the average across clans, weighted by the maximum number of retained ancestral amino acid sites across Pfams in each clan. We calculated standard errors (SE) using the weighted_se() function in the R package ‘diagis’^86^.

To assess relative usage, we divided the ancestral amino acid frequencies of pre-LACA, pre-LBCA clans, modern archaea or modern bacteria clans, by those of LUCA. We calculated Ses for usage using an approximation derived from a Taylor expansion of the ratio: 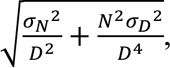 where *N* and *σ*_*N*_^2^ are the mean and variance of the numerator and *D* and *σ*_*D*_^2^ for the denominator^87^.

## Code and Data availability

All code and data relating to the analysis can be found on GitHub at https://github.com/sawsanwehbi/Ancient-Tryptophan-Tyrosine-usage.

## Author contributions

S.W. and J.M. conceived the study; A.W. inferred the archaea-trained time non-reversible amino acid substitution model, NQ.arch; S.W. performed research and analyzed data; N.L.T. and B.Q.M. contributed new analytic tools (developmental version of IQTREE3); D.S.L. revised manuscript; and S.W. and J.M. wrote the paper.

## Acknowledgements

We thank the Future Investigators in NASA Earth and Space Science and Technology (FINESST) program [80NSSC24K0384] and the Robert & Taylor Peterson Extraterrestrial Research Fund at the Arizona Astrobiology Center for funding S.W., the Chan-Zuckerberg Initiative [EOSS4-0000000312] for funding N.L.T. and B.Q.M., and the NSF [DEB-2333243] for funding A.W., B.Q.M., and J.M. We thank Jordan Douglas, Nigel Goldenfeld, Gavin Huttley, Aleksandar Radakovic, and Greg Fournier for helpful discussions, and Peter Goodman for writing the script for sequence downsampling.

