## Supplementary figures 1-8 and Supplementary Tables 1 and 2 for "Tryptophan became part of the universal genetic code post-LUCA"

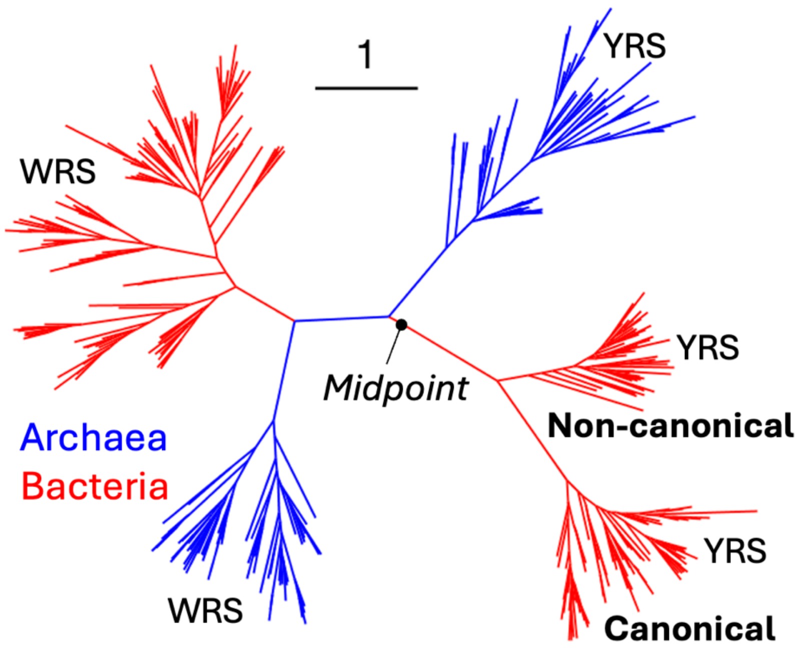


**Supplementary Figure 1. Bacterial YRS is deeply diverged into two groups: canonical and non-canonical.** The phylogeny is for PF00579 in prokaryotic WRS and YRS, as inferred by Wehbi et al.[8] but with all CPR bacteria and the truncated Euryarchaeota sequence 'G000403645' removed. Briefly, a maximum likelihood tree was reconstructed in IQ-Tree v2.1.3[32] and then reconciled with the WoL species tree[68] using GeneRax[88]. Branch lengths were then re-estimated with NQ.bac + R10. The tree, with sequences labelled by GenBank genome accession[89] in NCBI, is available on Github in 'PF00579_alnCPRdropped_NQbac.treefile'.

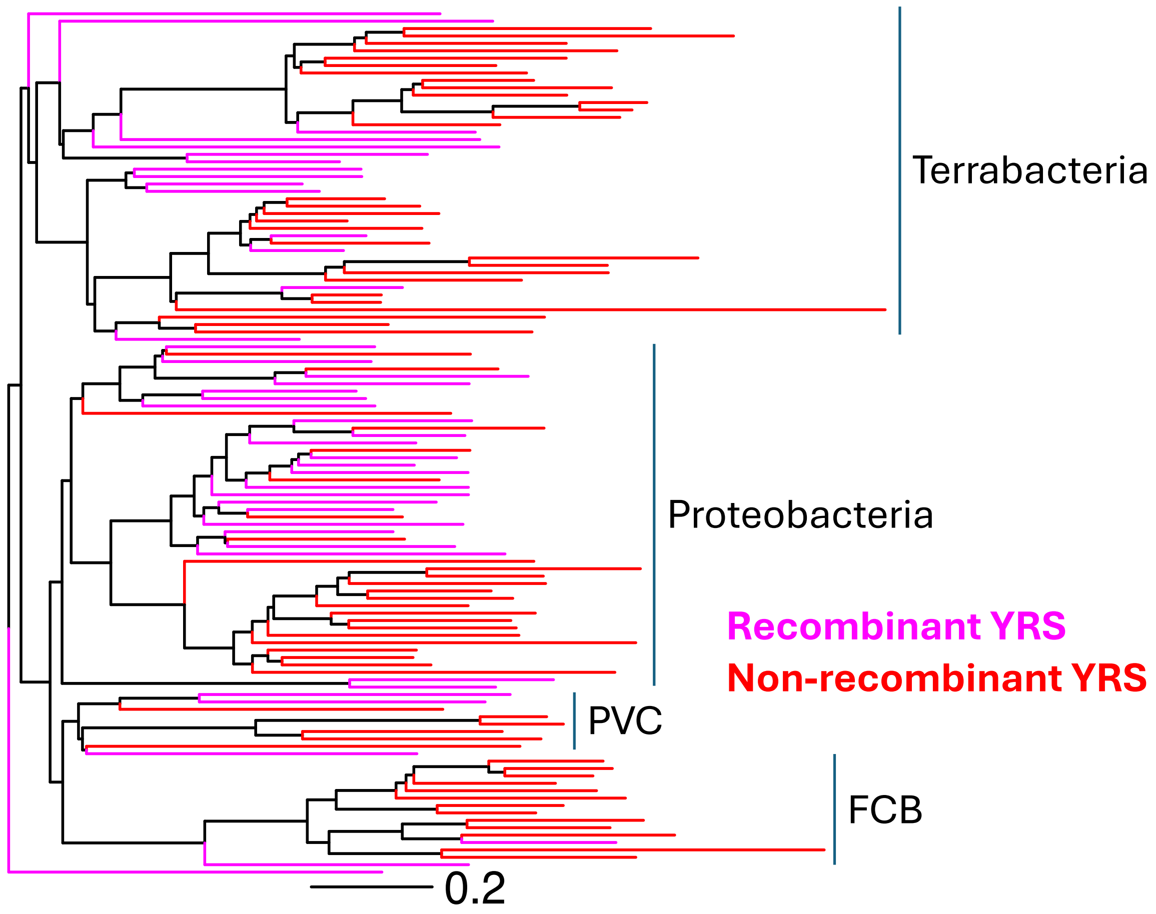


**Supplementary Figure 2. Distribution of recombinant and non-recombinant YRS in Bacteria.** Bacterial species tree and its branch lengths were adapted from the WoL microbial species tree, which used 381 marker genes, and an LG + G substitution model[68]. We retain only the bacterial species present in the YRS alignment shown in Figure 2A. (A). Pink and red branches denote species with the non-canonical and canonical YRS, respectively. The phylogenetic tree, with species labelled according to GenBank genome accessions[89] in NCBI, is in 'BacterialYRSspeciestree.newick'.

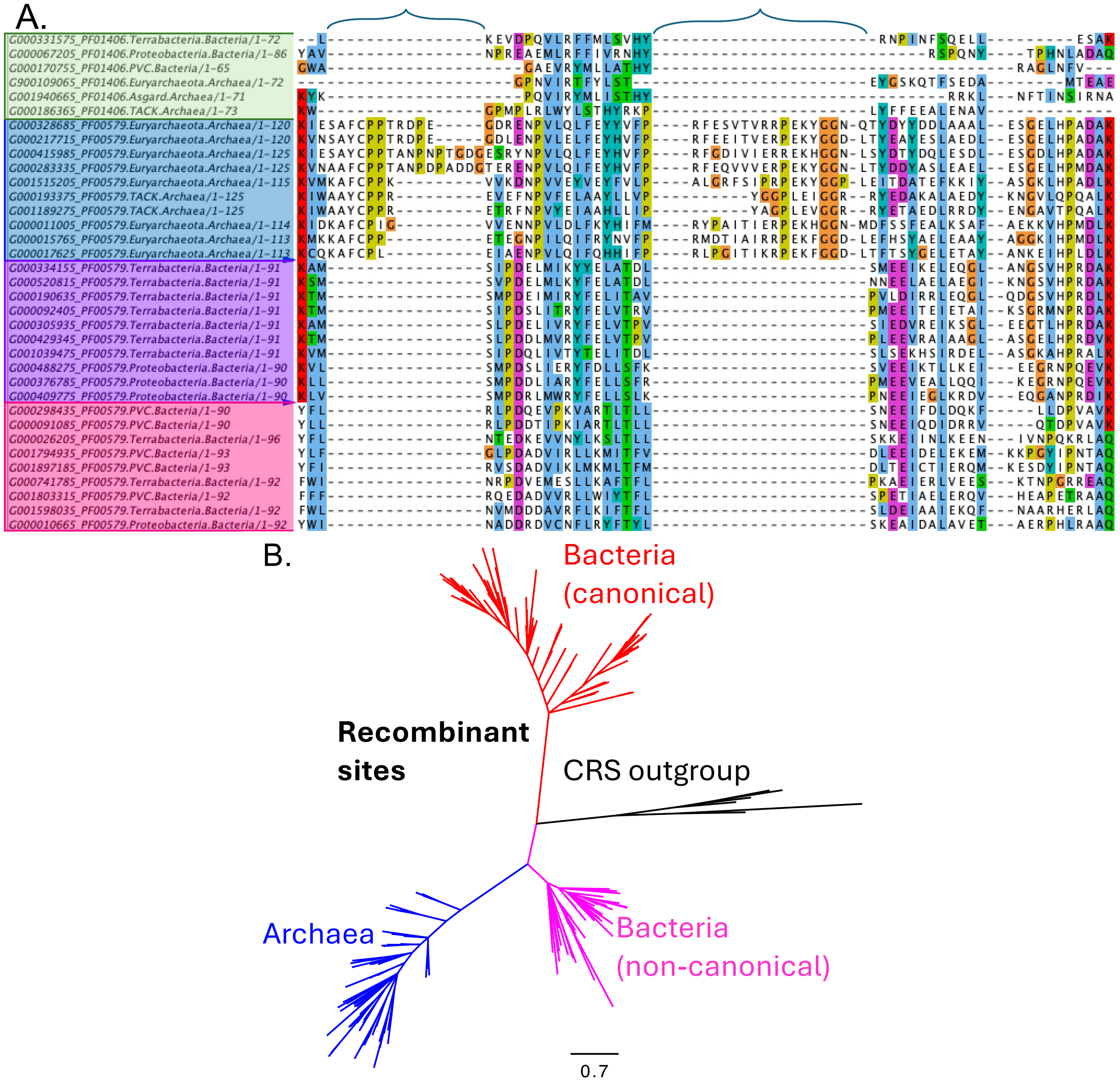


**Supplementary Figure 3. YRS recombinant sites analyzed together with a CRS outgroup.** A) Two regions that are present in archaeal YRS but absent in both bacterial YRS groups, are also absent in the CRS outgroup (indicated by blue curved brackets on top of the alignment), indicating that they evolved as insertions on the archaeal lineage. The first six sequences, highlighted in green, correspond to the CRS core catalytic domain (PF01406), the others are representatively downsampled from Figure 2A. B) Rooting with the CRS outgroup separates canonical bacterial from non-canonical bacterial and archaeal YRS. Tree was inferred from the putatively recombinant sites of the YRS alignment with six CRS outgroup sequences, three from each of Bacteria and Archaea. The tree topology was then inferred following the same process used to infer the trees in Figure 2B (see Methods for more details). The outgroup root is consistent with the branch-length based rooting methods (midpoint and MAD) in Figure 2B. Branch lengths were estimated using NQ.bac+R10. The YRS recombinant sites alignment with the PF01406 CRS outgroup is available on Github in 'YRS_alnPF01406_recombination.fasta'.

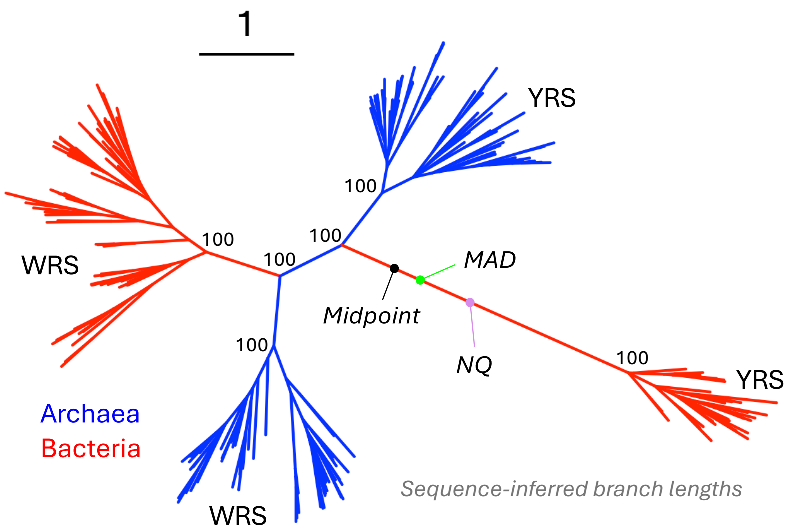


**Supplementary Figure 4. Using NQ.arch instead of NQ.bac+R10 yields the same roots as in Figure 3C, with greater confidence for midpoint and MAD.** Tree is available on GitHub in 'TruncPF00579_reconciledcon_arc.treefile'. With NQ.arch, the MAD root lies 26% and the midpoint root 17% of the way along from the archaeal split to the bacterial YRS radiation, compared to 23% and 10% with NQ.bac, respectively. The NQ root has 100% rootstrap support in both cases, with NQ.arch putting the root only 43% of the way toward the radiation of bacterial YRS, compared to 53% for NQ.bac.

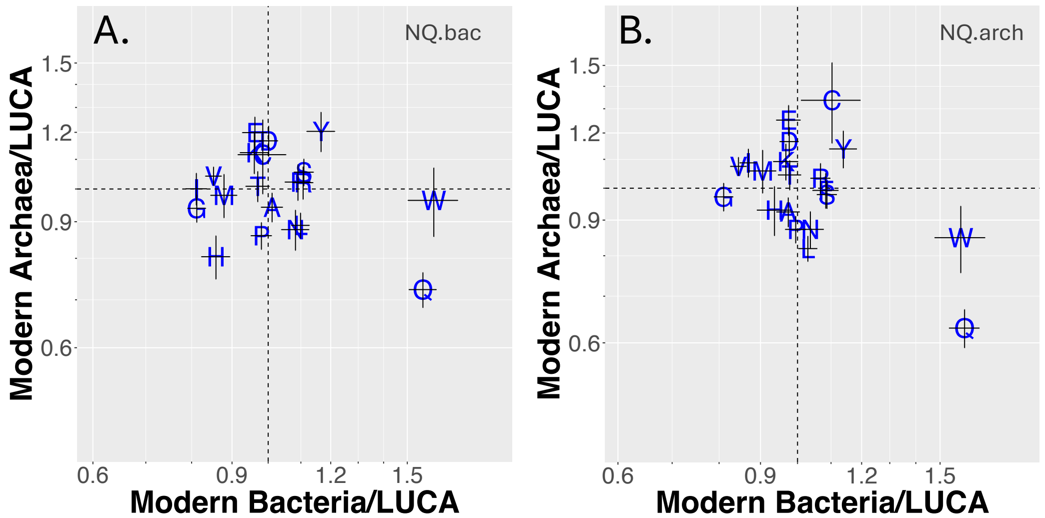


**Supplementary Figure 5. Recently emerged bacterial sequences are enriched for W relative to sequences dating back to LUCA, like the ancient bacterial sequences of Figure 4B.** See methods for identification of modern and LUCA sequence clans. Ratios of ancestral amino acid frequencies, reconstructed with either NQ.bac+R10 (A) or NQ.arch+R10 (B). Numerators used 163 modern archaea and 1578 modern bacteria clans, the denominator used 309 LUCA clans. Error bars indicate 66% confidence intervals (+/- 1 standard error; see Methods for details). Axes tick marks are drawn on a logscale, at intervals of 0.1.

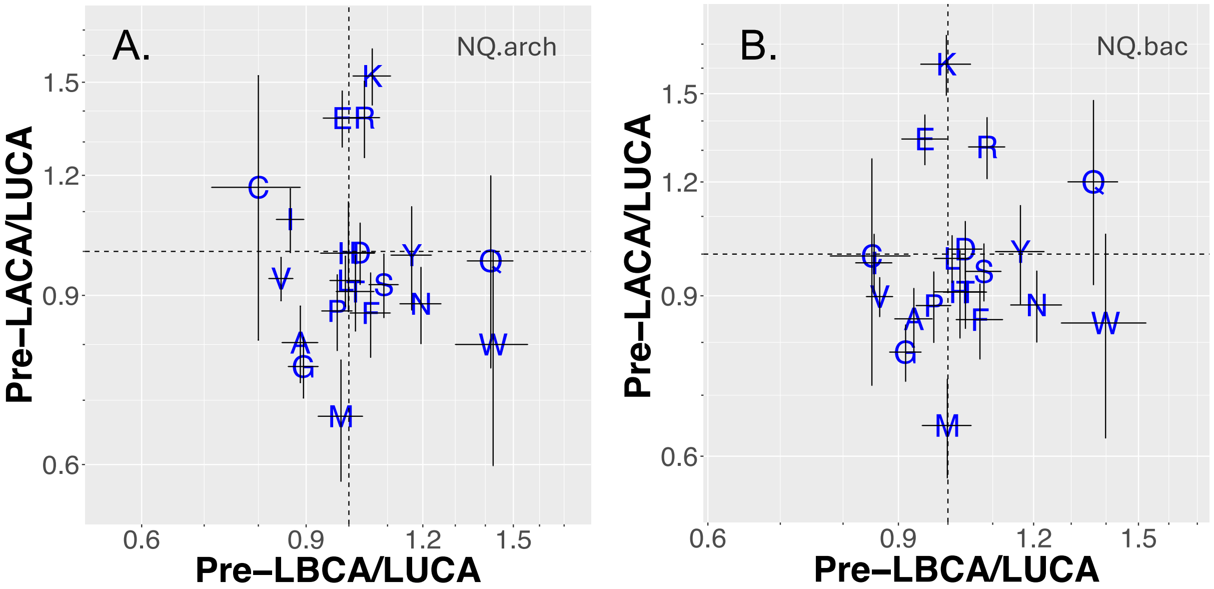


**Supplementary Figure 6. Higher W frequency in pre-LBCA than in pre-LACA or LUCA is also found using NQ.arch**. Panel B) with NQ.bac is identical Figure 4B, and is reproduced here to demonstrate insensitivity of amino acid usage ratios to model choice. Amino acid frequency reconstruction used either (A) NQ.arch+R10 or (B) NQ.bac+R10. Numerators used 31 pre-LACA and 251 pre-LBCA clans, the denominator use 309 LUCA clans. Error bars indicate 66% confidence intervals (+/- 1 standard error; see Methods for details). Axes tick marks are drawn on a logscale, at intervals of 0.1.

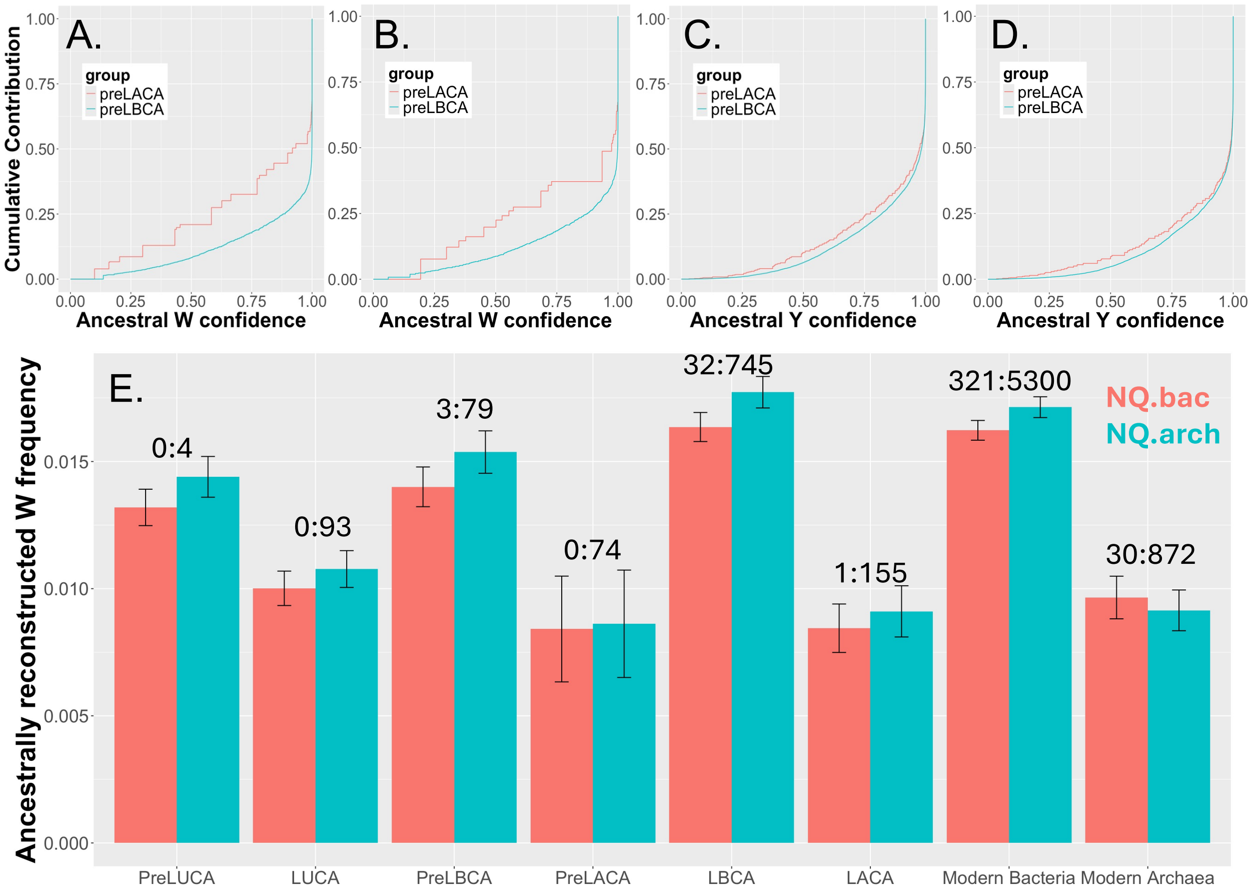


**Supplementary Figure 7. Absence of evidence for the presence of W in ancient archaea.** Inferred ancestral W in archaeal sequences comes from less confident sites than in bacterial sequences (A-B), in a manner not seen for Y (C-D). Ancestrally reconstructed W and Y frequencies, inferred with either NQ.bac (A, C) or NQ.arch (B, D), average the probability of W or Y across all reconstructed sites, broken down by confidence level on the x-axis. E) Columns show ancestrally reconstructed W frequency at sites that pass our quality filters, with NQ.bac+R10 or NQ.arch+R10. Ancestral W frequencies are slightly higher when inferred with NQ.arch than with NQ.bac. This can be explained by slower flux into and out of W in the NQ.arch model than in NQ.bact, use of which then keeps ancient W frequencies closer to contemporary frequencies; the normalized out-rates summed over the 19 amino acids, are 0.35 and 0.42, respectively. This difference between the models presumably reflects the fact that with fewer W residues in archaea, those few that are present are more highly conserved. For sequence ages shown on the x-axis, see Wehbi et al.[8] for annotation of LUCA and pre-LUCA clans, and the Methods of the current paper for annotation of pre-LBCA, pre-LACA, modern bacteria, and modern archaea clans. Numbers above the error bars indicate how many W : other amino acid sites are 100% "ultra-conserved" within the contemporary Pfam alignments from Wehbi et al.[8] whose age is given on the x-axis. Comparing the 32 Ws : 745 other ultra-conserved LBCA sites to only 1 W : 155 other ultra-conserved LACA sites yields p = 0.03 (Fisher's exact test). Results are in the same direction but statistically insignificant for pre-LBCA vs. pre-LACA. The main text reports on pooling these two groups.

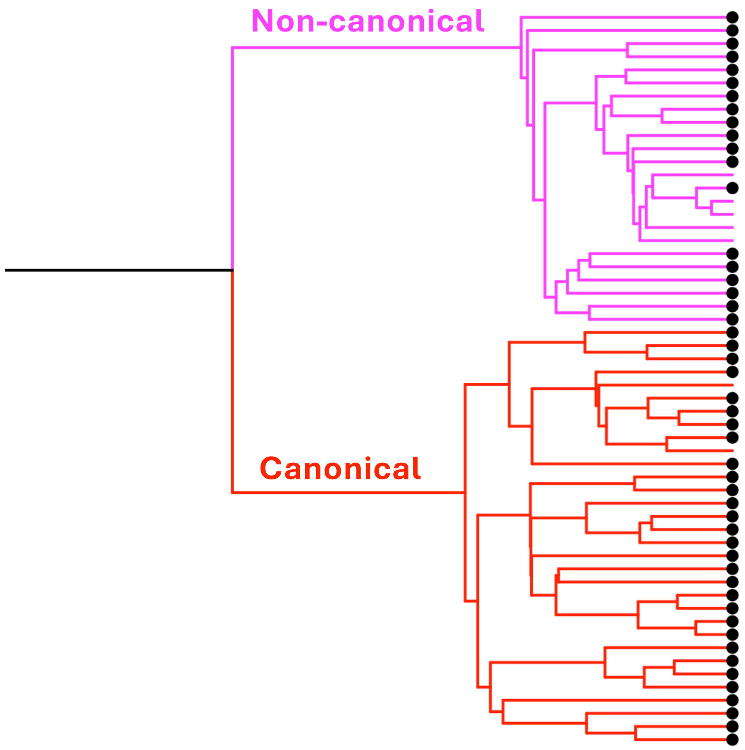


**Supplementary Figure 8. No strong phylogenetic pattern in the distribution of the additional S4 domain in bacterial YRS.** This is compatible with the high rates of horizontal gene transfer previously found for two highly diverged bacterial YRS groups[46]. We cross-referenced the YRS accessions with the Pfam database[90] on the Uniprot webserver[64] to match each protein with its associated Pfam domains. Out of 56 bacterial YRS proteins which were cross-referenced, UniProt annotated 49 as having the additional S4 domain – shown as a black circle at the tips. The phylogeny was extracted from the full WRS/YRS PF00579 inferred in Wehbi et al.[8] and pruned for the 56 YRS sequences for which UniProt provided information on the presence or absence of the S4 domain. We display the tree as ultrametric to artificially elongate very short branches, making the topology visible.

| Study | Rooting method | Data type | WRS origin |
|---|---|---|---|
| 1. Brown et al. 1997[11]<br>2. Yang et al. 2003[12] | Distance | Sequence | Pre-LUCA |
| 3. O'Donoghue and Luthey-Schulten 2003[14] | Outgroup | Structure | Pre-LUCA |
| 4. Nagel and Doolitle 1995[13] | Outgroup | Sequence | Pre-LUCA |
| 5. Ribas de Pouplana et al. 1996[18]<br>6. Patra et al. 2025[17] | Outgroup | Sequence | Post-LUCA |
| 7. Diaz-Lazcoz et al. 1998[15] | Other, non-outgroup | Sequence | Post-LUCA |
| 8. Dong et al. 2010[16] | Distance | Structure | Post-LUCA |

**Supplementary Table 1. Previous studies that rooted the YRS/WRS phylogenetic tree.** Some studies reached different conclusions despite using the same data type and rooting methods (e.g. Nagel and Doolittle 1995 versus Patra et al. 2025). In total, four studies support each of the two conclusions, emphasizing the need to revisit the WRS evolutionary history.

| Site in Alignment | Bacterial YRS MRCA | Archaeal YRS MRCA | WRS/YRS MRCA | Root | In YRS *C. pneumoniae* | In WRS *A. pernix* |
|---|---|---|---|---|---|---|
| *55* | A42 (0.73)<br>A42 (0.77) | L41 (0.54)<br>W41 (0.47),<br>L41 (0.42) | L 41(0.43),<br>W41 (0.27)<br>W41 (0.42),<br>L41 (0.32) | L41 (0.18),<br>A41 (0.16)<br>W41 (0.36),<br>L41 (0.27) | A42 | A41 |
| *127* | W96 (0.99)<br>W96 (0.99) | F83 (0.69)<br>F83 (0.60) | F83 (0.44),<br>L83 (0.19)<br>F83 (0.36),<br>L83 (0.24) | W83 (0.41),<br>F83 (0.23)<br>F83 (0.31),<br>L83 (0.21) | W90 | R84 |
| *249* | W168 (0.99)<br>W168 (0.99) | R149 (0.99)<br>R149 (0.99) | R153 (0.65)<br>R153 (0.50) | W155 (0.46),<br>R155 (0.29)<br>R155 (0.46),<br>K155 (0.29) | W162 | D149 |
| *606* | W211 (0.99)<br>W211 (0.99) | F195 (0.87)<br>F195 (0.88) | F197 (0.70)<br>F197 (0.72) | W199 (0.41),<br>F199 (0.30)<br>F199 (0.60) | W205 | F197 |
| *630* | W226 (0.99)<br>W226 (0.99) | Y211 (0.98)<br>Y211 (0.96) | Y213 (0.59)<br>Y214 (0.56) | W215 (0.41),<br>Y215 (0.30)<br>Y215 (0.50) | L220 | L213 |

**Supplementary Table 2. Ancestral sequence reconstruction reveals that only the bacterial YRS MRCA had W sites with high support (≥ 99%)**. All five sites at all four nodes of interest have <0.0002 probability of being an indel. For each of the five W sites, the most strongly supported amino acid(s) at the nodes of interest are shown with their probabilities inferred using NQ.bac (red), and NQ.arch (blue). Alignment coordinates refer to the PF00579 amino acid Muscle alignment available on GitHub in 'TruncPF00579_AA_Musclealn.fasta'. For ancestral sites with confidence < 50%, we also show the second most likely amino acid. We also give corresponding contemporary amino acids in YRS from the bacterium *Chlamydia pneumoniae* (G000091085) and WRS from the archaea *Aeropyrum pernix* (G000011125), illustrating less use of W in the latter.
